# HNF4 factors control chromatin accessibility and are redundantly required for maturation of the fetal intestine

**DOI:** 10.1101/610428

**Authors:** Lei Chen, Natalie H. Toke, Shirley Luo, Roshan P. Vasoya, Rohit Aita, Aditya Parthasarathy, Yu-Hwai Tsai, Jason R. Spence, Michael P. Verzi

## Abstract

As an embryo matures into a fetus, cells undergo remarkable transitions, accompanied by shifts in transcription factor regulatory networks and chromatin landscapes. The mechanisms of these developmental transitions are not completely understood. The embryonic intestine transitions from a rapidly proliferating tube with pseudostratified epithelium prior to embryonic day (E) 14.5, to an exquisitely folded columnar epithelium in the fetus. We sought to define factors that drive fetal maturation of the intestine. ATAC-seq profiling revealed a dramatic restructuring of intestinal chromatin during the embryonic-to-fetal transition, with CDX2 transcription factor motifs abundant at chromatin-accessible regions of the embryo, and hepatocyte nuclear factor 4 (HNF4) transcription factor motifs the most abundant in the fetal stages. Genetic inactivation of *Hnf4α* and its paralog, *Hnf4γ*, revealed that HNF4 factors are redundantly and vitally required for fetal maturation. In the embryo, CDX2 binds to and activates *Hnf4* gene loci to drive HNF4 expression at fetal stages. HNF4 and CDX2 transcription factors then occupy shared genomic regulatory sites and are required for chromatin accessibility at genes expressed in the maturing fetal intestine. Thus, intestinal transcription factor regulatory networks shift to accompany changing chromatin landscapes and gene expression profiles that occur during the transition of an embryonic tissue to its mature state.

## INTRODUCTION

The developing embryo is a collection of partially-fated cells that expand exponentially during the stages of organogenesis, approximately E9.5 to E13.5 in mice (Cao et al., 2019). As development proceeds into fetal stages, specified cells undergo transitions to acquire characteristics of mature tissues. The mechanisms of these developmental transitions are not completely understood, but are of great importance in understanding the basic mechanisms of development and developmental disorders, and in facilitating efforts of regenerative medicine.

The embryonic gut tube arises from endoderm after gastrulation and is specified along the anterior-posterior axis into distinct derivative endodermal organs. The primitive gut divides into foregut, midgut and hindgut; the small intestine develops primarily from the midgut. Murine gut tube formation is completed by E9.5, and its inner lining consists of a highly-proliferative, pseudostratified epithelium as the gut tube elongates, with the most rapid growth occurring by E14.5. At E14.5 to E15.5, the tissue undergoes a remarkable transition to a columnar epithelium and the process of villus formation, elongation, and maturation occurs at E15.5 to E18.5 (Chin et al., 2017; Walton et al., 2016a; Wells and Spence, 2014). The mechanisms triggering this embryonic to fetal transition in the developing gut remain unclear.

A shift in transcription factor regulatory networks accompanies the majority of known cellular transitions (Niwa, 2018; Wilkinson et al., 2017). Transcription factors function at distal genomic regulatory regions known as enhancers, which are the primary drivers of tissue-specific gene expression (Consortium, 2012; Nord et al., 2013; Shen et al., 2012; Visel et al., 2009). In our efforts to understand mechanisms driving developmental transitions in the gut, we recently mapped chromatin profiles of esophagus, forestomach, hindstomach and small intestine over developmental time. We noted a clear transition in chromatin accessibility within the developing intestine that corresponds to the stage in which the morphogenetic events reshaping the intestine occur (~E15.5). The transcription factor, CDX2, operates on both sides of this developmental transition. In the early embryonic intestine (prior to E13.5), CDX2 is required for intestinal specification, and loss of CDX2 leads to ectopic features of stomach and esophageal tissues in the intestine (Banerjee et al., 2018; Gao et al., 2009; Grainger et al., 2010; Kumar et al., 2019). Acute inactivation of CDX2 at later developmental timepoints (post-E13.5) compromises mature tissue functions, but intestinal identity is maintained (Banerjee et al., 2018; Kumar et al., 2019; Verzi et al., 2011). While CDX2 is a clear driver of intestinal specification and plays an additional role in driving fetal maturation, it remains unclear how this developmental transition occurs.

Here, we aim to find transcription factors that could function specifically in intestinal maturation. We find that genomic regions that become increasingly accessible at fetal stages are most enriched for DNA-binding motifs known to bind to the HNF4 transcription factor family. In adult intestine, HNF4A and HNF4G have been shown to drive expression of genes important for digestive physiology (Boyd et al., 2009; Cattin et al., 2009; Lindeboom et al., 2018). HNF4A binding activity is modulated by the microbiome (Davison et al., 2017) and suppressed in a mouse model of ulcerative colitis (Chahar et al., 2014). We recently demonstrated that HNF4A and HNF4G redundantly drive enterocyte identity in adult tissues (Chen et al., 2019). However, while HNF4 transcription factors have been studied in a number of developmental contexts, their redundant functions have not been assayed in the context of the developing gut.

*Hnf4α* deletion is embryonically lethal, with defects in visceral endoderm (VE) (Chen et al., 1994; Duncan et al., 1997). Complementation of *Hnf4α*^−/−^ embryos with *Hnf4α*^+/+^ VE by tetraploid aggregation (Duncan et al., 1997; Li et al., 2000) or conditional deletion of *Hnf4α* (Babeu et al., 2009; Garrison et al., 2006; Parviz et al., 2003) allowed for the investigation of HNF4 function in later developmental stages. In the liver, *Hnf4α* was dispensable for liver specification but was essential for hepatocyte differentiation (Hayhurst et al., 2001; Li et al., 2000; Parviz et al., 2003). However, no consequence of HNF4A loss in the developing small intestinal epithelium has been observed. This may be due to an inefficient depletion in the small intestine due to the mosaic action of the *Foxa3-Cre* driver, as suggested by Garrison *et al*. (Garrison et al., 2006). Inactivation of conditional *Hnf4α* alleles using the *Villin-Cre* driver resulted in no embryonic phenotype (Babeu et al., 2009), possibly due to the relatively late onset of *Villin-Cre* expression (Madison et al., 2002) or unappreciated genetic redundancy with *Hnf4γ*. Here, we demonstrate that intestinal expression of the *Hnf4* paralogs is induced during the embryonic to fetal transition, and that *Hnf4* genes depend upon CDX2 for expression in both mice and humans. Analysis of chromatin accessibility and ChIP-seq data in the developing and adult gut supports a model in which the transcription factor CDX2 activates *Hnf4* gene expression, and together HNF4 and CDX2 drive tissue maturation by activating genes important for intestinal function in post-natal life. Finally, we generate a mouse model lacking HNF4 factors in the embryonic intestine to show that while HNF4 factors are dispensable for intestinal specification and villus morphogenesis, they are critical for maturation of the fetal intestine.

## RESULTS

### Chromatin landscapes indicate that HNF4 factors are more likely to function in intestinal maturation than specification

Deciphering the mechanisms of tissue maturation is a critical goal in understanding developmental disorders and facilitating efforts in regenerative medicine. Developmental transitions are accompanied by changes in chromatin accessibility and transcription factor regulatory networks. To better understand the regulatory mechanisms driving intestinal maturation, we re-analyzed ATAC-seq data (Banerjee et al., 2018) to identify chromatin-accessible regions in isolated intestinal epithelial cells (Fig.1A). Three categories of accessible regions were identified (MACS *P* value ≤ 10^−5^). 2,644 regions (cluster 1) were accessible at E11.5 and remained accessible at all stages examined. 10,544 regions (cluster *2*, “maturation enriched regions”) of intestinal chromatin are progressively accessible, whereas 30,702 regions (cluster 3, “embryo enriched regions”) lose accessibility from E11.5 to adult (Fig.1A, Table S1). Notably, a gain in accessibility at “maturation enriched regions” coincides with the loss of accessibility at “embryo enriched regions”. This E14.5-16.5 stage is marked by a transition of the intestinal epithelium from a pseudostratified to columnar tissue. To identify the regulatory complexes likely operating during this developmental transition, we applied DNA-binding motif analysis. Regions selectively accessible in the maturing gut exhibit HNF4A/G as the top-scoring motif (Fig.1B, Table S1), suggesting that HNF4 factors function in maturation of the developing gut. Conversely, CDX2 is the most prevalent transcription factor motif in the 30,702 regions that are more accessible in embryonic epithelium (Fig.1B, Table S1). H3K27ac (Kazakevych et al., 2017), which marks active enhancer regions, is enriched at stage-specific accessible chromatin regions defined by ATAC-seq (Fig.1C), corroborating putative regulatory function of these regions. Gene ontology analysis suggests that genes nearby the accessible regions of the early embryonic intestine function in intestinal specification and morphogenesis, whereas genes nearby accessible chromatin in the maturing intestine exhibit function consistent with the mature tissue (Fig.1D, Table S1). DNA-binding motif enrichment analysis at each timepoint revealed that while CDX2 binding motifs are enriched as early as E11.5, HNF4 binding motifs are first detected at E14.5 and become more prevalent at accessible chromatin regions of fetal and adult stages (Fig.S1A). Mirroring their motif enrichment profiles, we find that *Cdx2* transcripts are highly and equally expressed across early embryonic to adult stages, whereas *Hnf4* transcripts are not robustly expressed until fetal and adult stages (Fig. 1E). Protein expression levels of HNF4 factors (Fig.S1B,C) appear to increase over developmental time, consistent with the increase in transcript levels. Similarly, human *CDX2* expression is observed upon intestinal specification of hESC-derived endoderm by FGF and WNT, whereas expression of *HNF4A* and *HNF4G* is not observed until cultures are further matured into human intestinal organoids (Fig. 1F, Fig.S1D). Taken together, the chromatin landscapes, DNA motif enrichment, and expression profiles of intestinal epithelial cells across developmental time suggest that HNF4 factors are more likely to function in tissue maturation than intestinal specification.

**Fig. 1.**
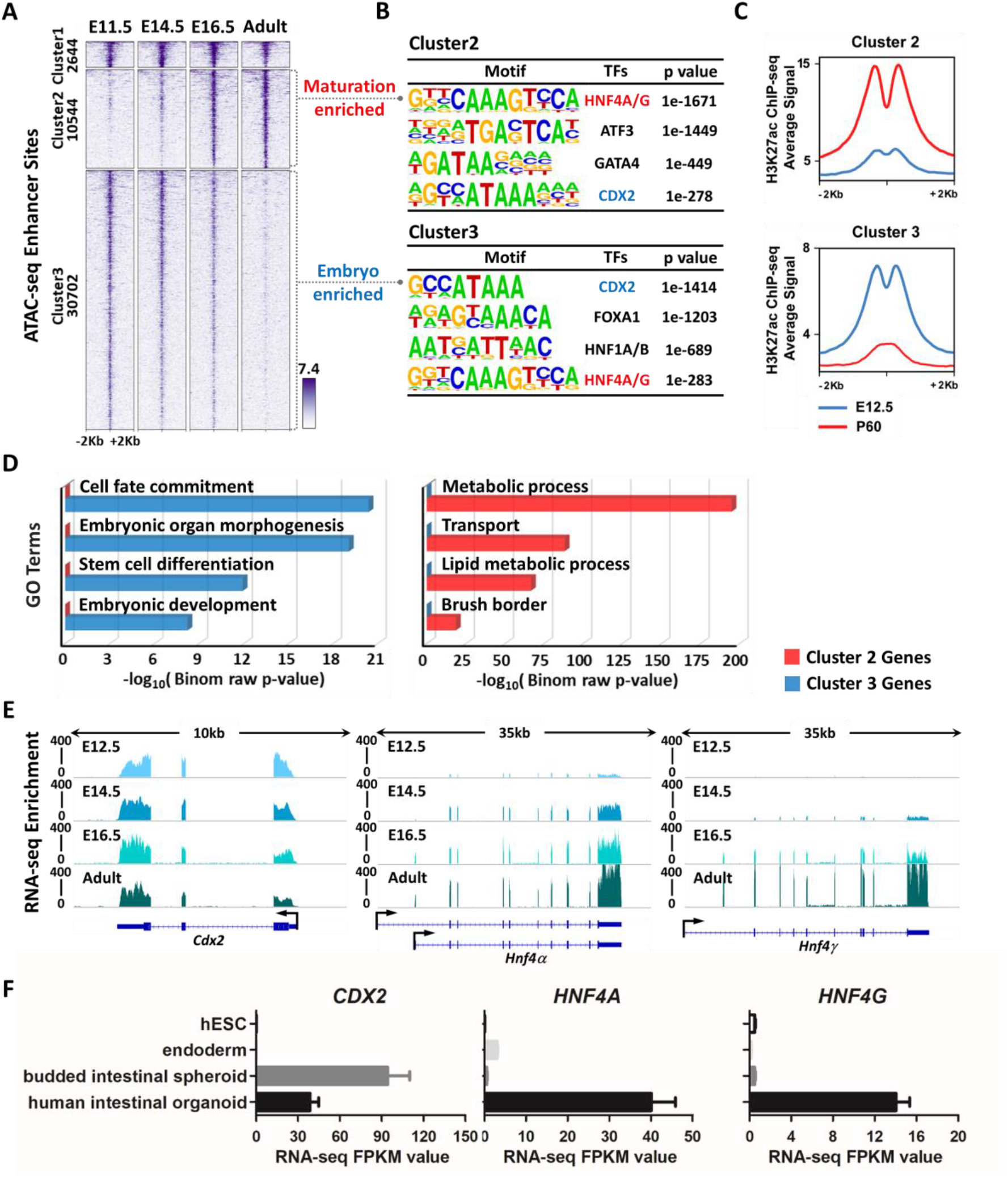
Chromatin landscapes indicate that HNF4 factors function in maturation of the developing gut. (A) ATAC-seq (GSE115541, n = 2 biological replicates per timepoint, isolated embryonic epithelium was collected from the entire small intestine) defined regions of accessible chromatin across indicated stages of mouse intestinal epithelium development (MACS *P* value ≤ 10^−5^). *K*-means clustering analysis reveals a shift in accessible enhancer chromatin of intestinal epithelial cells across developmental time. (B) HOMER *de novo* DNA-motif enrichment analysis of ATAC-seq regions (MACS *P* value ≤ 10^−5^) shows that CDX2 binding sequences are more prevalent in accessible regions of the early embryonic state (embryo enriched, cluster 3), whereas HNF4 binding sequences are more prevalent in accessible regions of the fetal and adult states (maturation enriched, cluster 2). (C) H3K27ac ChIP-seq (GSE89684, n = 2 biological replicates per timepoint) profiles demonstrate that the active chromatin marker is enriched in a stage-specific manner, corresponding to embryo-specific or P60 adult-specific accessible regions. (D) GREAT GO term analysis shows distinct gene ontologies of target genes linked to stage-specific ATAC-seq sites within 20kb. (E) RNA-seq of purified epithelium (GSE115541, n = 2 biological replicates per timepoint, isolated embryonic epithelium was collected from the entire small intestine) shows that *Cdx2* is highly and equally expressed across developmental time, whereas *Hnf4α* and *Hnf4γ* are not robustly expressed until E14.5 and E16.5, respectively. (F) RNA-seq (E-MTAB-4168) shows that *CDX2* expression is induced when human endoderm is specified to intestine by treatment with FGF4 (500 ng/ml) and CHIR99021 (2 μM, WNT agonist). By contrast, *HNF4A/G* expression is not induced until human intestinal organoids are formed by subsequent differentiation steps. See schematic in Fig.S1D for details on human intestinal organoid differentiation conditions. Data are presented as mean ± SEM. At least 3 biological replicates are included per stage, and samples from the same stage are grouped and presented.

### HNF4 factors are dispensable for intestinal specification and villus morphogenesis, and CDX2 functions upstream of HNF4 factors

To test the function of HNF4 in the developing gut, we deleted a conditional allele of *Hnf4α* (Hayhurst et al., 2001) in the gut endoderm using the *Shh-Cre* driver (Harfe et al., 2004), which is activated in the intestinal epithelium starting at ~E9.5. We also examined germline null embryos lacking *Hnf4γ* mediated by CRISPR knockout (Chen et al., 2019), and embryos with both HNF4 paralogs simultaneously deleted (hereafter referred to *Hnf4α^KO^, Hnf4γ^KO^*, and *Hnf4αγ^DKO^*, Fig.S1E,F). As we and others have reported (Banerjee et al., 2018; Gao et al., 2009; Grainger et al., 2010; Kumar et al., 2019), *Cdx2^KO^* in the early embryonic gut epithelium leads to an intestine exhibiting hindstomach characteristics (ATPase and foveolar PAS positive cells) in the jejunum and stratified squamous esophageal characteristics (p63 positive cells) in the ileum. However, similar characteristics are not observed in the *Hnf4αγ^DKO^* embryos (Fig.2A,B), suggesting that HNF4 factors are dispensable for intestinal specification. These findings are consistent with the analysis of accessible chromatin (see above, Fig. 1), in which CDX2, but not HNF4, motifs are enriched at regulatory regions most active in the embryonic intestine (Fig.1B, Fig.S1A).

**Fig. 2.**
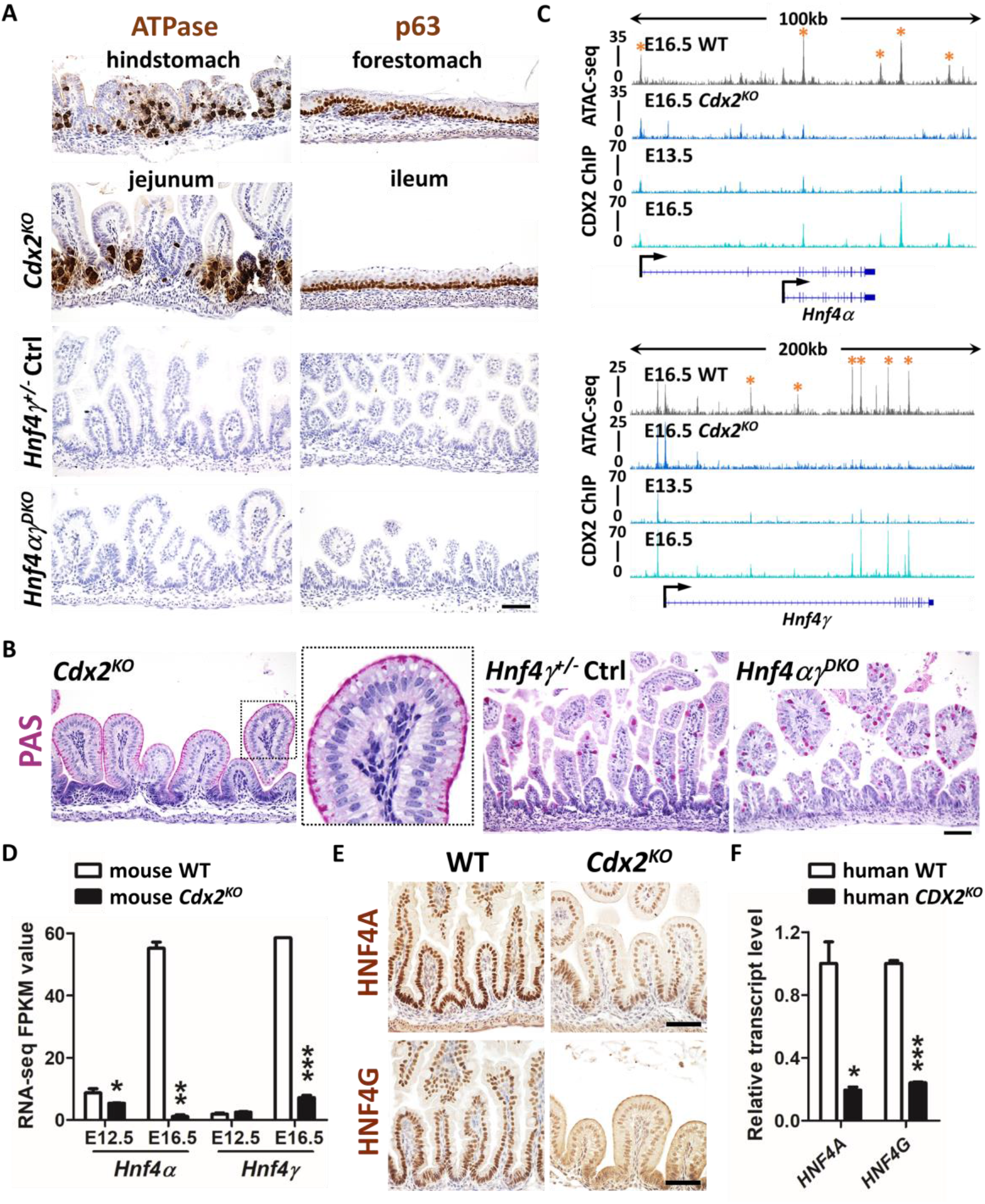
HNF4 is activated by CDX2 in the developing gut, but HNF4 is not required for intestinal specification. (A) Immunostaining of ATPase and p63 (representative of 4 biological replicates) in E18.5 control, *Shh-Cre;Cdx2^f/f^* (*Cdx2^KO^*) and *Shh-Cre;Hnf4α^f/f^;Hnf4γ^Crispr/Crispr^* (*Hnf4αγ^DKO^*) embryos. The hindstomach marker ATPase is ectopically expressed in the *Cdx2^KO^* jejunum but not in the control and *Hnf4αγ^DKO^* jejunum. The forestomach marker p63 is ectopically expressed in the *Cdx2^KO^* ileum (squamous mucosa) but not in the control and *Hnf4αγ^DKO^* ileum. (B) PAS staining indicates that normal intestinal goblet cells are replaced with cells resembling gastric foveolar cells (PAS positive cells at apical cell surface) in the *Cdx2^KO^* jejunum of E18.5 embryos,but not in the control or *Hnf4αγ^DKO^* jejunum (representative of 4 biological replicates). (C) CDX2 binds to *Hnf4α* and *Hnf4γ* loci at E13.5 and E16.5 (CDX2 ChIP, GSE115314, n = 2 biological replicates, whole small intestine epithelium), and accessible chromatin is compromised at gene loci of *Hnf4α* and *Hnf4γ* upon CDX2 loss (ATAC-seq, GSE115314, n = 2 biological replicates, whole small intestine epithelium). Asterisks denote putative regulatory regions. (D) The transcript levels of *Hnf4α* and *Hnf4γ* are significantly downregulated in the intestinal epithelial cells of the *Shh-Cre;Cdx2^KO^*. Data are presented as mean ± SEM (RNA-seq, GSE115541, n = 2-3 biological replicates, whole small intesitne epithelium). Statistical tests are embedded in DESeq2 at *P* < 0.001***, *P* < 0.01** and *P* < 0.05*). (E) Immunostaining of HNF4A and HNF4G shows reduced protein levels of HNF4 paralogs in E18.5 *Cdx2^KO^* (representative of 4 biological replicates). Scale bars, 50 μm. (F) qRT-PCR shows reduced transcript levels of *HNF4A* and *HNF4G* in human intestinal organoids derived from *CDX2^CrisprKO^* hES cells compared to organoids derived from control cells. Data are presented as mean ± SEM (n = 3 biological replicates, Student’s t-test, two-sided at P < 0.001*** and *P* < 0.05*).

Since CDX2 is required for intestinal specification but HNF4 factors appear dispensable, we hypothesized that CDX2 may function upstream of HNF4 factors. Examination of CDX2 ChIP-seq revealed that CDX2 binds to loci of *Hnf4α* and *Hnf4γ* at E13.5 and the ChIP-seq signal strengthens at E16.5 (ChIP-seq panel in Fig.2C). Loss of CDX2 results in loss of the chromatin accessibility at the gene loci of *Hnf4* factors (ATAC-seq panel in Fig.2C) and reduced *Hnf4* transcript (Fig.2D) and protein levels (Fig.2E), suggesting that CDX2 is a direct activator of *Hnf4α*. *Hnf4γ* expression is similarly dependent upon CDX2 (Fig.2C-E); although due to the later onset of *Hnf4γ* expression, these differences first become apparent at E16.5 (Fig.2D). In developing human intestine, HNF4 factor expression is similarly dependent upon CDX2, as reflected by measurement of *HNF4* transcript levels in human intestinal organoids derived from control or *CDX2^CrisprKO^* hESCs (Fig.2F). These results indicate that expression of HNF4 factors is dependent upon CDX2 during fetal maturation of the intestine.

CDX2 binds to both maturation enriched and embryo enriched regions of intestinal accessible chromatin (ATAC-seq regions from Fig.1A) at E13.5. When the intestine is mature, the CDX2 and HNF4 ChIP-seq signal is most robust at maturation enriched regions, with comparatively less binding at embryonic-accessible regions (Fig.3A,B), suggesting that these factors work to activate maturation-specific gene expression. Consistent with this idea, genes such as *Myo7b* and *Pls1* (Fig.3C), which are bound by both HNF4 and CDX2, exhibit increased transcript expression during tissue maturation. To test whether HNF4 and CDX2 function is required for chromatin accessibility at regions that become increasingly accessible in the fetal stages, we compared ATAC-seq signal in control and mutant embryos. Chromatin accessibility is indeed compromised at maturation enriched regions from the intestinal epithelium of both E16.5 *Cdx2^KO^* and *Hnf4αγ^DKO^* (Fig.4A-E). Chromatin accessibility loss in these mutants is selective to maturation-enriched chromatin, as chromatin accessiblity at promoters is unaltered, and serves as an internal control. Taken together, these data suggest an active role of CDX2 and HNF4 factors in binding and maintaining accessibility at maturation-accessible regions.

**Fig. 3.**
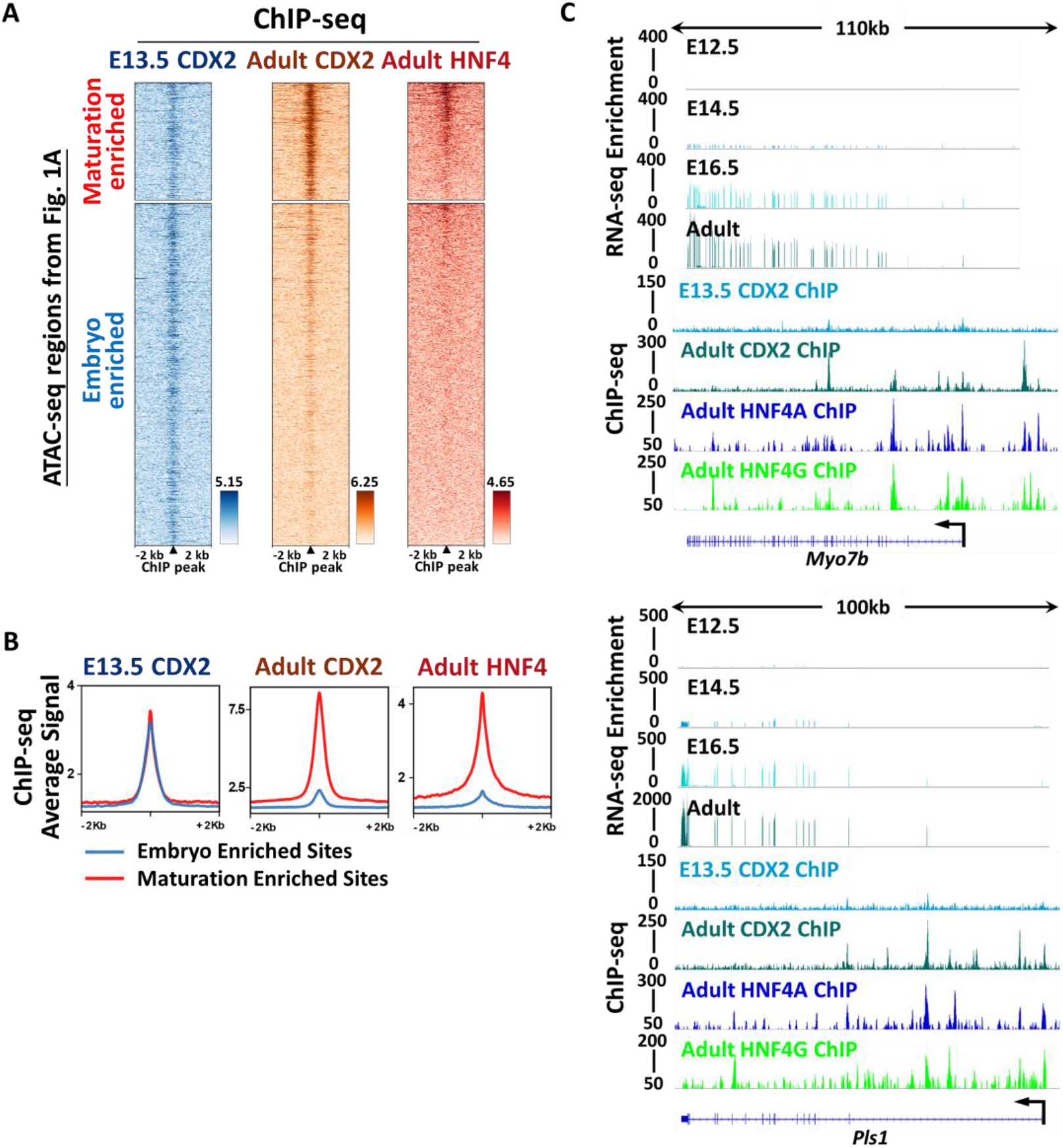
Chromatin regions that become accessible in the fetal tissue are bound by CDX2 and HNF4. Genes nearby these regions are activated at fetal stages. (A-B) ChIP-seq profiles show that CDX2 (GSE115314, n = 2 biological replicates, whole small intestine epithelium) binds to both maturation enriched (cluster 2 of Fig.1A) and embryo enriched (cluster 3 of Fig.1A) regions of intestinal accessible enhancer chromatin in E13.5 embryos. When tissues become mature, CDX2 (GSE34568 and GSE115314, n = 2 biological replicates) and HNF4 (GSE112946, n = 2 biological replicates per HNF4A and HNF4G ChIP, adult duodenum epithelium) bind more robustly to the maturation enriched regions rather than to the embryo enriched regions. The peaks of CDX2 ChIP in the heatmaps are aligned to the binding events of HNF4 ChIP. (C) Examples of genes located at maturation enriched regions are visualized using IGV. Brush border genes, such as *Myo7b* and *Pls1*, are expressed when maturation occurs at E16.5 and are robustly expressed when intestines become mature in the adult (RNA-seq panels, GSE115541, n = 2 biological replicates). These genes are directly bound by CDX2 (GSE34568 and GSE115314, n = 2 biological replicates) and HNF4 (GSE112946, n = 2 biological replicates per HNF4A and HNF4G ChIP).

**Fig. 4.**
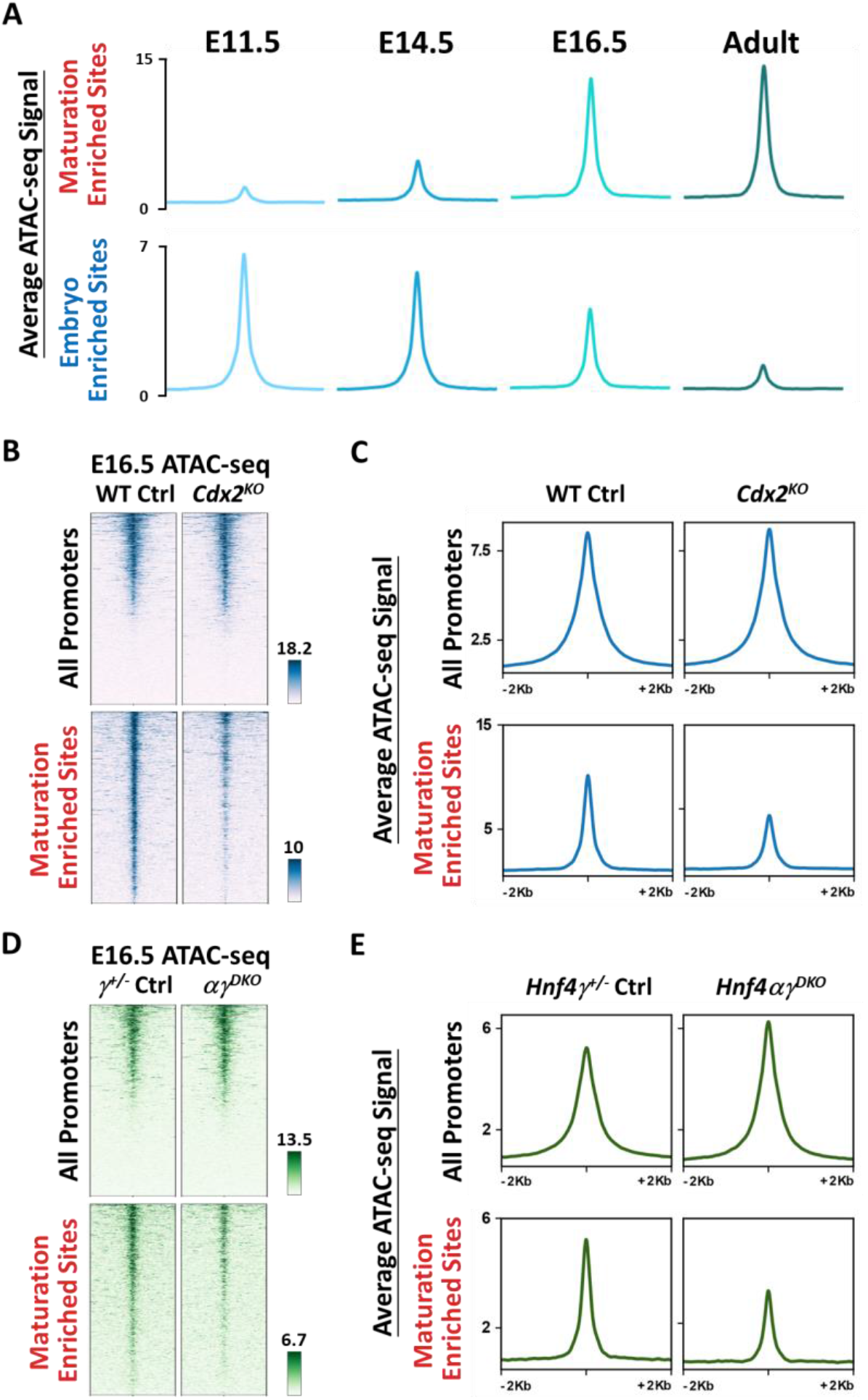
Maturation-enriched regions of accessible enhancer chromatin show loss of accessibility in the intestinal epithelium of *Cdx2^KO^* and *Hnf4αγ^DKO^* mutants. (A) The intestinal chromatin (GSE115541, n = 2 biological replicates) becomes more accessible at maturation enriched regions and less accessible at embryo enriched regions (ATAC-seq regions defined from Fig.1A) from E11.5 embryo to adult. The intestinal chromatin accessibility at the maturation enriched sites is compromised upon depletion (*Shh-Cre*) of *Cdx2* (B-C, GSE115314) or *Hnf4* (D-E) in E16.5 embryos, whereas promoter regions are relatively unaffected and serve as an internal control (n = 2 WT controls, 1 *Hnf4γ*^+/−^ controls, 2 *Cdx2^KO^* mutants and 2 *Hnf4αγ^DKO^* mutants, whole small intestine epithelium).

HNF4 factors thus appear dispensable for tissue specification, but their expression pattern and control of maturation-enriched regulatory regions suggest potential roles during later developmental stages. From E14.5 to E15.5, the pseudostratified intestinal epithelium resolves to a cuboidal epithelium, and mesenchymal cells crosstalk with epithelial cells to initiate villus formation (Chin et al., 2017; Walton et al., 2016a; Wells and Spence, 2014). For example, PDGFRα-expressing mesenchymal clusters serve as signaling centers for villus morphogeneis (Walton et al., 2012). We investigated whether villus morphogenesis is disrupted upon HNF4 loss. At E15.5, *Hnf4αγ^DKO^* intestines are similar to their littermate controls (Fig.5A), and there are no obvious defects in epithelial morphology, mesenchymal condensations (PDGFRα positive cells), or proliferation (BrdU positive cells) upon HNF4 loss at E14.5 (Fig.5B). These results suggest that proliferation and tissue morphogenesis initiate properly in the absence of HNF4 paralogs. Consistent with the observation of normal villus morphogenesis in HNF4 mutants, adult HNF4 bound-regions (defined by ChIP-seq, Chen et al., 2019) are not very accessible during embryonic stages, but become markedly more accessible following villus morphogenesis (Fig.5C,D). Chromatin accessibility at promoter regions serves as an internal control for each timepoint (Fig.5C,D). Genes linked to HNF4-ChIP-seq regions also exhibit increased expression after villus morphogenesis, as the tissue matures (RNA-seq, Fig.5E). Together, these findings suggest that HNF4 likely functions after villus morphogenesis to drive gene expression during the maturation of the developing fetal gut.

**Fig. 5.**
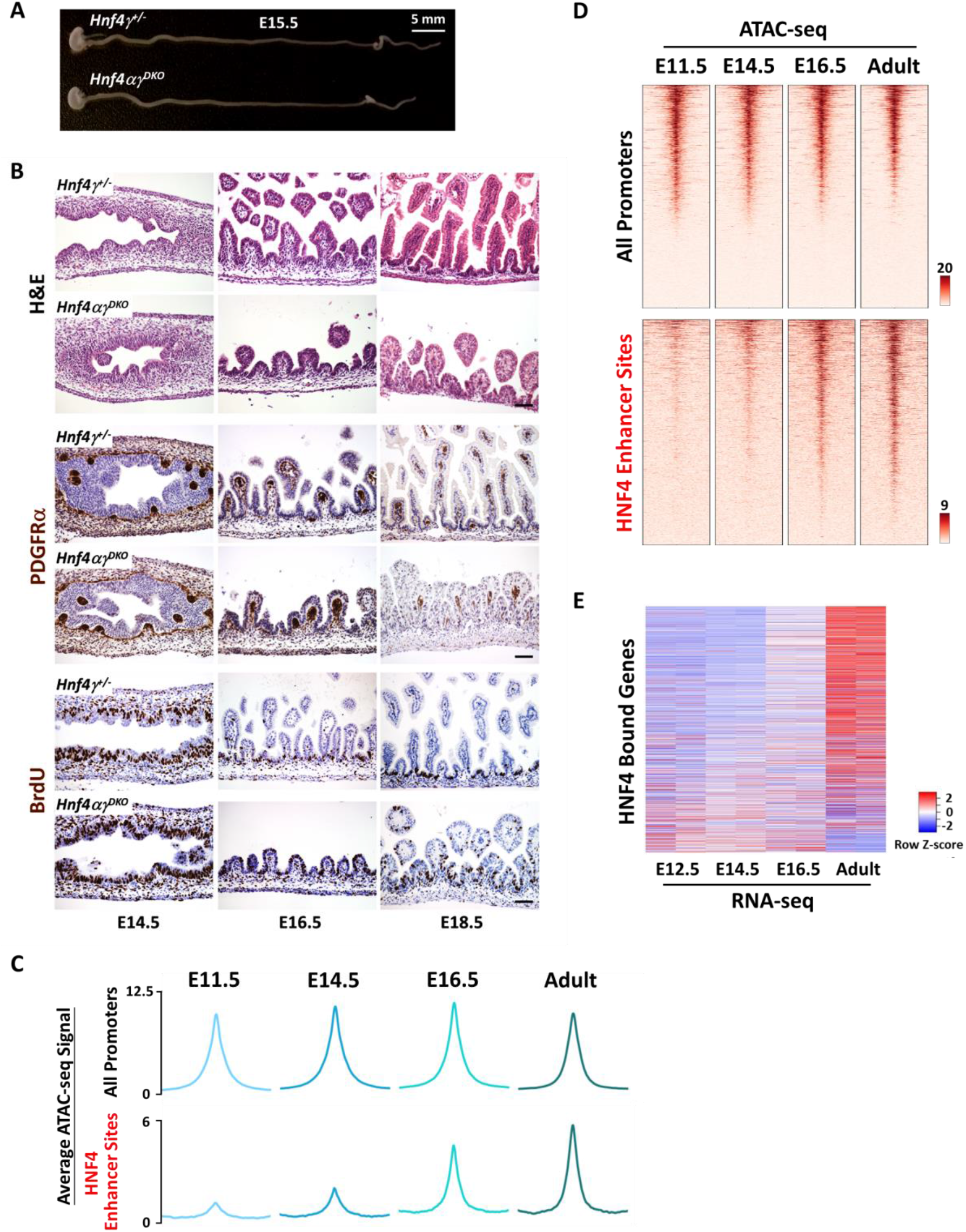
Villus morphogenesis initiates despite the loss of HNF4, and chromatin accessibility and gene regulation at HNF4-binding sites suggest a role for HNF4 after villus morphogenesis. (A) Whole mount images of representative intestines dissected from *Hnf4αγ^DKO^* and littermate *Hnf4γ*^+/−^ control embryos at E15.5 (n = 4 biological replicates per genotype). (B) H&E staining and immunostaining of PDGFR*α* and BrdU across developmental time show no striking differences, indicating that HNF4 factors are dispensable for the onset of villus morphogenesis (representative of 4 biological replicates). The pregnant female mice were injected with 1 mg BrdU 1 h before euthanasia. Scale bars, 50 μm. (C-D) Chromatin becomes accessible during fetal and adult stages at enhancer regions bound by HNF4 in the adult epithelium (MACS *P* value ≤ 10^−3^, GSE112946, n = 2 biological replicates per ChIP), indicating a potential role for HNF4 in activating genes nearby these regions during intestinal maturation. (E) Increased transcript levels of HNF4-bound genes in intestinal epithelial cells are observed across developmental time from E12.5 to adult villi. Genes with transcriptional start sites within 20 kb of HNF4 enhancer bound sites were used for plotting a heatmap showing the relative transcript expression levels over the indicated developmental stages (GSE115541, n = 2 biological replicates per timepoint).

### HNF4 factors are redundantly required for fetal maturation of the intestine

From E15.5 to E18.5, villi elongate into the lumen and cells that leave the intervillus regions exit the cell cycle and begin to express markers of differentiation. To better appreciate the function of HNF4 factors during this timepoint, we compared ATAC-seq on E16.5 intestinal epithelial cells isolated from *Hnf4αγ^DKO^* and control embryos. 5,391 accessible chromatin regions lose accessibility upon HNF4 loss (cluster 2 in Fig.6A, Table S2). Genes nearby these HNF4-dependent regions are associated with mature intestinal functions, such as adhesion, brush border formation, and lipid metabolism (Fig.6B, Table S2). For example, chromatin accessibility increases at regions of the brush border gene *Enpep* as the tissue matures (Fig.6C), and accessibility is almost completely lost in the *Hnf4αγ^DKO^* epithelium (Fig.6D). HNF4 paralogs directly bind to many maturation-specific genes (ChIP-seq in Fig.6E and Fig.S2), and transcript levels of these maturation-specific genes are dramatically reduced in *Hnf4αγ^DKO^* (Fig.6F).

**Fig. 6.**
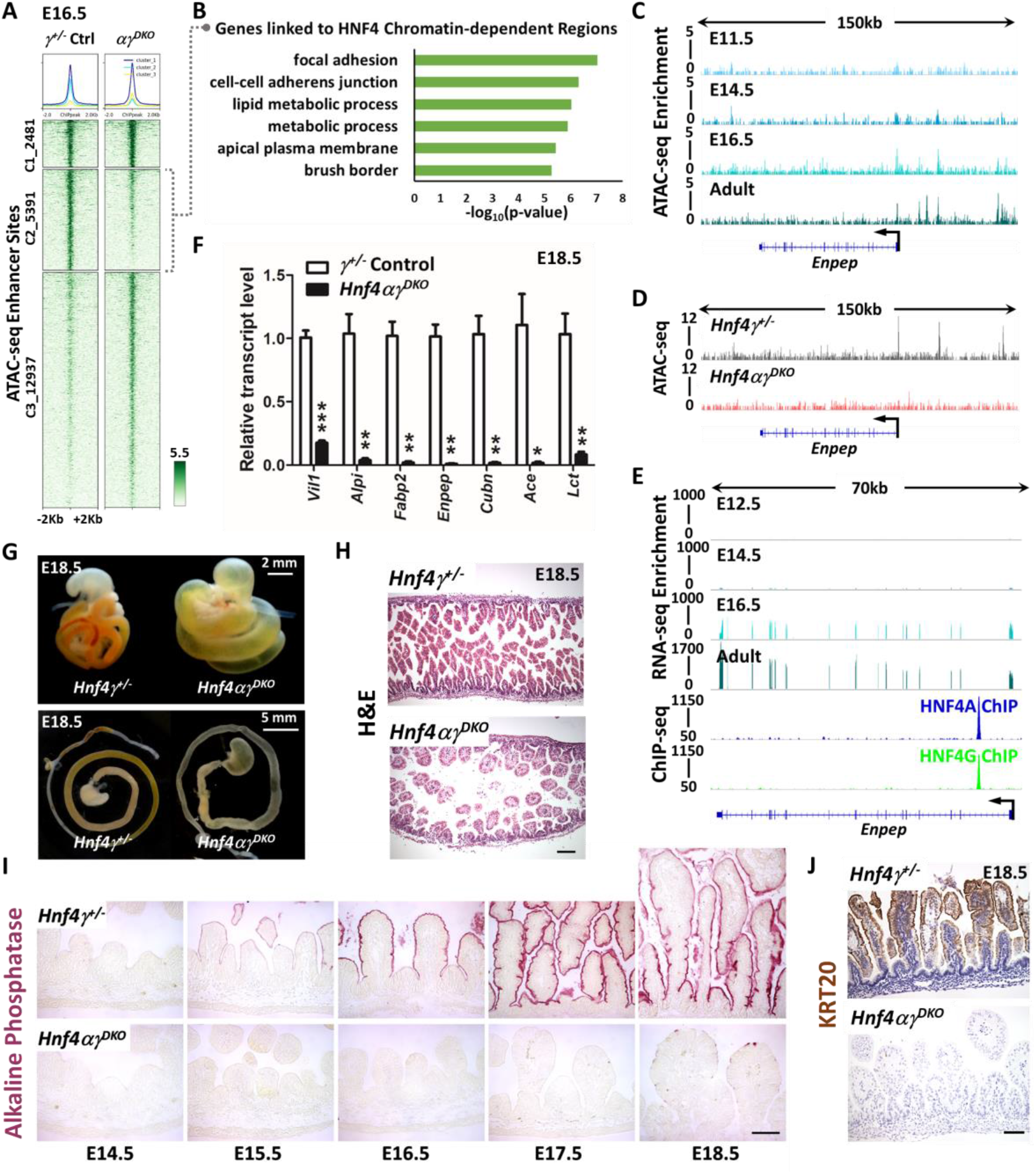
HNF4 paralogs are redundantly required for fetal maturation of the intestine. (A) *K*-means clustering of ATAC-seq data collected from E16.5 intestinal epithelial cells isolated from *Hnf4αγ^DKO^* and control embryos identify 5,391 regions (cluster 2) that are dependent on HNF4 factors for chromatin accessibility (n = 2 biological *Hnf4αγ^DKO^* replicates and 1 *Hnf4γ*^+/−^ control in this study; MACS *P* value ≤ 10^−5^). (B) Functional annotation of the genes linked to HNF4-dependent accessible chromatin by DAVID. Genes with transcriptional start sites within 20 kb of ATAC-seq sites of cluster 2 (loss of chromatin accessibility upon HNF4 loss) from panel 6A were used for analysis. (C-E) Examples of chromatin accessibility at HNF4 dependent ATAC-seq sites are visualized using IGV. (C) ATAC-seq (GSE115541) shows a time-dependent increase of chromatin accessibility at locus of the brush border gene *Enpep* from E11.5 embryo to adult. (D) ATAC-seq shows compromised chromatin accessibility at the *Enpep* locus is observed in E16.5 *Hnf4αγ^DKO^*. (E) RNA-seq (GSE115541) shows corresponding increase in transcript levels of *Enpep* over developmental time, and HNF4A and HNF4G directly bind to *Enpep* (GSE112946, ChIP-seq panel), suggesting a direct regulation. (F) qPCR shows that the transcript levels of genes known to be expressed in the mature intestine are dramatically reduced in *Hnf4αγ^DKO^* compared to that of the littermate *Hnf4γ*^+/−^ controls in isolated E18.5 intestinal epithelial cells. Data are presented as mean ± SEM (n = 4 controls and 5 mutants, Student’s t-test, two-sided at P < 0.001***, *P* < 0.01** and *P* < 0.05*). (G) Whole mount images of E18.5 intestine indicate that loss of HNF4 paralogs leads to an underdeveloped intestine with distended and translucent lumen (representative of 6 biological replicates). (H) Strikingly stunted villi are observed in E18.5 *Hnf4αγ^DKO^* embryos compared to littermate *Hnf4γ*^+/−^ controls, as revealed by H&E staining (representative of 4 biological replicates, E18.5 duodenum; see expanded panel in Fig.S3D). (I) *Hnf4αγ^DKO^* exhibits diminished alkaline phosphatase staining (differentiation marker) across developmental time (representative of 4 biological replicates, E14.5-E18.5 duodenum). (J) Keratin 20 immunostaining also indicates compromised intestinal differentiation in *Hnf4αγ^DKO^* embryos (representative of 4 biological replicates, E18.5 duodenum). Scale bars, 50 μm. Also see Fig.S2 and Fig.S3 for additional metrics of intestinal maturation.

Consistent with the loss of expression of maturation genes, *Hnf4αγ^DKO^* intestines exhibit a translucent and distended lumen at E18.5, suggesting an underdeveloped intestine compared to their littermate controls (Fig.6G). The gross morphological defects in intestines are more severe with loss of 3 or 4 *Hnf4* alleles compared to loss of 1 or 2 *Hnf4* alleles (Fig.S3A). After birth, *Hnf4αγ^DKO^* pups are not able to survive, whereas *Hnf4α^KO^γ*^+/−^ pups can survive but exhibit growth retardation (Fig.S3B,C), which may be due to incomplete intestinal maturation. At E18.5, villi are noticeably shorter in mutants lacking 3 or 4 *Hnf4* alleles but not in the *Hnf4* single mutants or controls (Fig.6H, Fig.S3D), suggesting that HNF4 paralogs are redundantly required for villus elongation and extension into the developing gut lumen. There is no obvious increase in apoptosis in *Hnf4αγ^DKO^*, as evaluated by cleaved caspase 3 staining (Fig.S4A). The differentiation marker, alkaline phosphatase, which is localized to the apical surface of villus enterocytes, is normally expressed in the controls beginning at E15.5 in the maturing tissue and increases over developmental time (top panels in Fig.6I). However, alkaline phosphatase is not detectable in *Hnf4* double mutants (bottom panels in Fig.6I). As expected for genetic redundancy between *Hnf4* paralogs, *Hnf4* single mutants show normal expression of alkaline phosphatase (Fig.S4B). Proliferative cells, which are normally restricted to the intervillus regions of the fetal gut, are expanded into the villi of *Hnf4αγ^DKO^* (Fig.S4C), which may be attributed to the loss of villus differentiation in mutants lacking HNF4 factors (Fig.6I,J). Taken together, our results indicate that HNF4 paralogs are dispensable for specification and villus morphogenesis in the developing gut, but are redundantly required for fetal maturation through direct binding to fetal maturation genes.

## DISCUSSION

The regulatory mechanisms governing the transition of embryonic to mature tissue is a significant frontier for both developmental biology and regenerative medicine. Chromatin accessibility dynamics across intestinal development provide new insights into the fundamental molecular basis of intestinal specification and maturation. HNF4 motifs are most prevalent in the accessible chromatin during the maturation of the developing gut, and we provide evidence that HNF4 transcription factors are indeed important for maturation of the fetal intestine. Here, we build a model that CDX2 functions in gut specification, binds to and activates HNF4 factors in the developing gut. Together, these factors are required to mature the fetal tissue, ultimately achieving a stabilized and mature intestine.

Interestingly, lower-level CDX2 binding in the embryo is observed at regions that will become accessible in fetal stages. (Fig.3A). Lower level CDX2 ChIP-signal at these poorly accessible regions could reflect a “low-level sampling” behavior that has recently been described for the FOXA1 transcription factor (Donaghey et al., 2018). When ectopically expressed in fibroblasts, FOXA1 exhibits low ChIP-seq signal at binding sites typically exclusive to other cell lineages, such as liver or endoderm, a behavior that could relate to FOXA1’s relatively strong interaction with chromatin and slow nuclear mobility compared to other transcription factors (Sekiya et al., 2009). Low-level FOXA1 binding is strengthened at these sites by co-expression of partner factor GATA4. HNF4 factors could similarly be stabilizing CDX2 at maturation-specific regions. A potential partner for CDX2 at embryo-specific regions remains elusive. It will be interesting to test whether CDX2 exhibits similar nuclear mobility as FOXA1.

It is also interesting to note that CDX2 appears to function in both the embryonic and fetal transcription factor regulatory networks (Fig.7). Transitions between regulatory networks could more seamlessly occur when certain factors are present across the transition, such as the presence of *Sox2* and *Esrrb* in both embryonic stem cells and trophoblast stem cells (Adachi et al., 2013). Rather than to shut down an entire set of transcription factors and establish an entirely new set of factors, the common presence of CDX2 might function as a placeholder to transition from the embryonic to fetal networks. Inhibitory factors have also been shown to play a role in tissue maturation, such as *Blimp1* (Harper et al., 2011; Mould et al., 2015).

**Fig. 7.**
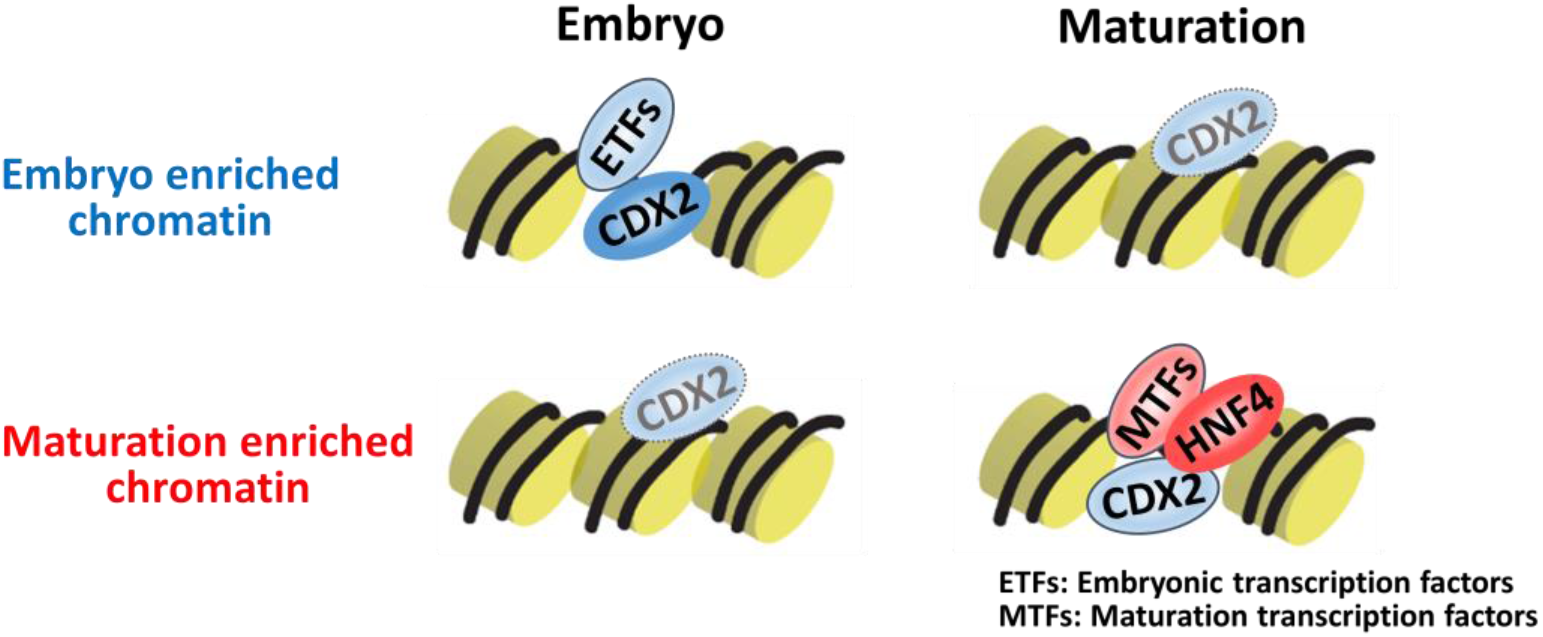
Potential model of how intestinal transcription factor networks shift as embryos mature to adults. Left, CDX2 functions in gut specification, and robustly binds to embryo enriched accessible chromatin regions with other embryonic transcription factors. In the embryo, lower-level CDX2 binding may also occur at regions that will become accessible in the fetus. Right, as intestine matures to fetal stages, maturation-enriched regions become more accessible. A new transcription factor network, highlighted by HNF4 works together to stabilize enhancer chromatin and drive intestinal maturation.

Transcription factors and developmental signaling pathways function in complex and collaborative networks to promote proper tissue development and function. BMP signals control intestinal villus patterning (Walton et al., 2016b) and intestinal looping (Nerurkar et al., 2017), but the functions and mechanisms of BMP signals in villus maturation are not fully explored. In a recent study, we identify a reinforcing feed-forward loop of BMP/SMAD signaling and HNF4, which promotes and stabilizes enterocyte cell identity in the adult intestine (Chen et al., 2019). Future studies can also investigate the presence of this reinforcing loop of BMP/SMAD signaling and HNF4 in the developing gut, and whether it promotes the maturation of the developing intestine or stabilization of mature transcription factor networks. These findings will build on the field’s knowledge of intestinal development and may help influence efforts of regenerative medicine to provide healthy intestinal tissue to patient populations.

## MATERIALS AND METHODS

### Mice

The *Shh-Cre* transgene (Harfe et al., 2004), *Cdx2^f/f^* (Verzi et al., 2010), *Hnf4α^f/f^* (Hayhurst et al., 2001), and *Hnf4γ^Crispr/Crispr^* (Chen et al., 2019) alleles were bred to achieve the indicated genotypes. The *Shh-Cre*^−^ embryos served as littermate controls unless otherwise indicated. The day on which a vaginal plug was found was considered to be E0.5. Embryonic tail biopsies were used for rapid genotyping using KAPA Mouse Genotyping Kits (Kapa Biosystems, KK7352). All mouse protocols and experiments were approved by the Rutgers Institutional Animal Care and Use Committee. All samples were collected between 12:00 and 14:00 to avoid circadian variability.

### Histology and Immunostaining

Intestinal tissues were fixed with 4% paraformaldehyde at 4°C overnight, washed with PBS, and dehydrated through ascending alcohols prior to paraffin embedding. 5 μm thick paraffin sections were used for histological staining. Hematoxylin (VWR, 95057-858) and eosin (Sigma, HT110180) staining was performed using standard procedures. Alkaline phosphatase activity was detected using the AP Staining Kit II (Stemgent). For Periodic Acid-Schiff (PAS) staining, slides were incubated in 0.5% periodic acid and stained with Schiff’s Reagent (Alfa Aesar, J612171). In the case of tissue prepared for BrdU immunohistochemistry, the pregnant female mice were injected with 1 mg BrdU, and embryos were harvested 1h after injection. Immunohistochemistry was performed using primary antibodies against HNF4A (Santa Cruz sc-6556 X, 1:2000), HNF4G (Santa Cruz sc-6558 X, 1:2000), ATPase (MBL International Corp. D032-3, 1:200), P63 (Santa Cruz sc-8343, 1:500), BrdU (Bio-Rad MCA2060, 1:500), Ki67 (Abcam ab16667, 1:300), PDGFRα (Santa Cruz sc-338, 1:1000), Cleaved Caspase 3 (Cell Signaling 9661, 1:200), and Keratin 20 (Cell Signaling 13063, 1:2500). After incubating with secondary antibody and the Vectastain ABC HRP Kit (Vector Labs), slides were developed using 0.05% DAB (Amresco 0430) and 0.015% hydrogen peroxide in 0.1 M Tris. After mounting, the slides were viewed on a Nikon Eclipse E800 microscope. Images were photographed with a Retiga 1300 CCD (QImaging) camera using QCapture imaging software. When adjustments of sharpness, contrast, or brightness were made, they were applied uniformly for comparative images.

### Human embryonic stem cell (hESC) culture and differentiation

All hESCs work was reviewed and approved by the University of Michigan human pluripotent stem cell research oversight committee (HPSCRO). The hESC cell line H9 (WA09, NIH stem registry #0062) was obtained from the WiCell Research Institute. *CDX2^CrisprKO^* hESCs were generated as described previously (Kumar et al., 2019). hESCs were maintained and differentiated into endoderm, hind/midgut and human intestinal organoids as published previously (Kumar et al., 2019; Tsai et al., 2017).

### Intestinal epithelial cell isolation

Embryos were collected at E18.5, and the freshly harvested embryonic small intestine (caudal stomach to rostral caecum) was opened longitudinally with forceps, cut into 1 cm pieces, and then rotated in 3 mM EDTA in PBS at 4 °C for 40 min. To release the epithelial cells from underlying muscular tissue, the tissue was vigorously shaken after EDTA incubation. The supernatant was collected as the whole epithelium fraction. Cells were pelleted by centrifugation at 170 g at 4 °C and then washed with cold PBS, and processed for RNA extraction using Trizol (Invitrogen, 15596018) according to the manufacturer’s protocols.

### RNA extraction and qRT-PCR

The RNA was reverse transcribed using SuperScript III First-Strand Synthesis SuperMix (Invitrogen, 18080-400) with Oligo(dT)_20_ primers to prepare cDNA. qRT-PCR analysis was performed using gene-specific primers and SYBR Green PCR Master Mix (Applied Biosystems, 4309155). The sequences of the primers used are available upon request. The 2^−ΔΔCt^ method was applied to calculate the fold change of relative transcript levels, and *Hprt1* was used for normalization.

### Single intestinal epithelial cell isolation for ATAC-seq

Embryos were collected at E16.5, and the freshly harvested embryonic small intestine (caudal stomach to rostral caecum) was opened longitudinally with forceps and cut into 1 cm pieces. Intestinal tissues were treated with pre-warmed 0.25% Trypsin for 8 min at 37°C on a vortex station (speed set between 6-7), neutralized with 10% FBS, and passed through a 70-μm cell strainer. Cells were stained with the anti-CD326 (EpCAM) magnetic microbeads antibody (Miltenyi Biotec, 130-105-958) for 40 min on ice. To obtain single magnetic antibody conjugated EpCAM positive epithelial cells, stained cells were passed through a 40-μm cell strainer and then collected over a MS column (Miltenyi Biotec, 130-042-201) in a magnetic field. 20,000 cells were used for ATAC-seq as described previously (Buenrostro et al., 2013; Buenrostro et al., 2015) with slight modifications. Briefly, cells were resuspended in ice-cold lysis buffer (10 mM Tris, pH 7.4, 10 mM NaCl, 3 mM MgCl2, and 0.1% NP-40), and then centrifuged at 500 g for 10 min at 4°C. The isolated nuclei were incubated with Nextera Tn5 Transposase (Illumina FC-121-1030) at 37°C for 30 min. The transposed chromatin was purified with QIAquick PCR Purification Kit (QIAGEN), and PCR was amplified with high-fidelity 2X PCR Master Mix (New England Biolabs M0541). One-third of the maximum fluorescent intensity was used to determine the additional cycles. The PCR amplified libraries were purified, and fragment size was selected using Pippin Prep and sequenced on Illumina NextSeq 550.

### Bioinformatics analysis

For ATAC-seq and ChIP-seq analysis, raw sequencing reads (fastq) were quality checked using fastQC (v0.11.3) and were further aligned to mouse (mm9) genomes using bowtie2 (v2.2.6) (Langmead and Salzberg, 2012) to obtain bam files. Deeptools bamCoverage (Ramirez et al., 2016) (v2.4.2, duplicate reads ignored, RPKM normalized and extended reads) was used to generate bigwig files from bam files, and BigWigMerge (v2) was used to merge the bigwig files of different replicates. MACS (v1.4.1) (Zhang et al., 2008) was used for peak calling and to generate bed files from bam files, and BEDTools (v2.17.0) (Quinlan, 2014) was used to merge, intersect or subtract the intervals of bed files. Promoters were defined within 2 kb of the transcription start sites of RefSeq genes, and enhancers were defined by excluding promoters. Haystack (v0.4.0) (Pinello et al., 2018) quantile normalized bigwigs were used to create *k*-means clustering heatmaps of ATAC-seq using computeMatrix and plotHeatmap from deeptools (v2.4.2) (Ramirez et al., 2016). Genomic regions of desired *k*-means clusters were extracted from bed files generated by plotHeatmap for further analysis. Homer findMotifsGenome.pl (v4.8.3, homer *de novo* Results) (Heinz et al., 2010) was used to identify transcription factor motifs enriched at peaks. Genes associated with peaks were identified using the peak2gene/BETA-minus function (v1.0.2) in Cistrome tools (Liu et al., 2011). Enriched gene ontologies were identified from genomic regions (bed file) using GREAT analysis (v3.0.0) (McLean et al., 2010) or DAVID (v6.8) (Huang da et al., 2009). SitePro (v1.0.2) (Shin et al., 2009) was used to visualize the average signals of ChIP-seq in the desired genomic regions. The Integrative Genomics Viewer (IGV 2.4.13) (Robinson et al., 2011) was used to visualize bigwig tracks.

For RNA-seq analysis, raw sequencing reads (fastq) were quality checked using fastQC (v0.11.3), and were further aligned to mouse (mm9) genomes using Tophat2 (v2.1.0) (Kim et al., 2013) to generate bam files. Kallisto (v0.44.0) (Bray et al., 2016) was utilized to quantify the transcript abundances of the RNA-Seq samples through *pseudoalignment*, using single-end reads and an Ensembl mm9 transcriptome build index. Then, the tximport (v1.8.0) (Soneson et al., 2015) package was run in R (v3.5.2) to create gene-level count matrices for use with DESeq2 (v1.20) (Love et al., 2014) by importing quantification data obtained from Kallisto. DESeq2 was then used to generate RPKM values per kilobase of gene length per million mapped fragments at each time course point with comparison of *Cdx2^KO^* replicates and wild-type replicates. DESeq2 was also used to generate p-values of gene matches. Genes with FPKM>1, a commonly used minimal expression threshold, were used for further analysis. Heatmapper (Babicki et al., 2016) was used to display relative transcript levels of genes of interest by using normalized RPKM values. The Integrative Genomics Viewer (IGV 2.4.13) (Robinson et al., 2011) was used to visualize bam tracks.

### Statistical analysis

The data is presented as mean ± SEM, and statistical comparisons were performed using two-sided Student’s t-test at *P* < 0.001***, *P* < 0.01** or *P* < 0.05*. Bioinformatics related statistical analysis was done with the embedded statistics in each package, including HOMER (Heinz et al., 2010), GREAT (McLean et al., 2010), DAVID (Huang da et al., 2009) and DESeq2 (Love et al., 2014). *P* < 0.05 (95% confidence interval) was considered statistically significant.

## Acknowledgements

We thank Madhurima Saxena and Ramesh Shivdasani for helpful suggestions and curating published data. We thank Kenneth Zaret for helpful discussions.

## Author contributions

L.C. conceived and designed the study, performed benchwork and bioinformatics, collected and analyzed the data, and wrote the manuscript; N.H.T., S.L., R.P.V. and A.P. contributed to benchwork. R.A. contributed to bioinformatics; J.R.S and Y-H.T. provided human intestinal organoid data; M.P.V. conceived, designed and supervised the study, and wrote the manuscript.

## Data availability

All ATAC-seq data of this study have been deposited in GEO (GSE128674). The following secure token has been created to allow review of record GSE128674: cpgbwsyarrghpeh. The following datasets from GEO were reanalyzed: The accession numbers for the transcriptome and chromatin accessibility of time-course WT and *Cdx2^KO^* from our previous studies are GSE115314 (Kumar et al., 2019) and GSE115541 (Banerjee et al., 2018). The accession numbers for the CDX2 ChIP-seq and HNF4 ChIP-seq from our previous studies are GSE34568 (Verzi et al., 2013), GSE115314 (Kumar et al., 2019) and GSE112946 (Chen et al., 2019). GSE89684 (Kazakevych et al., 2017) was used to mark active chromatin with the time-course H3K27ac ChIP-seq. The accession number for the RNA-seq data of differentiation of hESCs into human intestinal organoids is E-MTAB-4168 (Tsai et al., 2017) in the ArrayExpress database.

## Supplementary information

**Table S1**. Genome coordinates for ATAC-seq performed in intestinal epithelial cells from E11.5 embryo to adult, including 30,702 embryonic enhancer regions (cluster 3 from Fig.1A) and 10,544 maturation enhancer regions (cluster 2 from Fig.1A). Additionally, the full list of HOMER *de novo* motif-calling analysis on these embryonic and maturation enriched regions are reported respectively. Finally, the results of GO term enrichment using GREAT analysis for genes with their transcriptional start sites within 20 kb of these embryonic and maturation regions are reported respectively. These data correspond to findings in Fig.1.

**Table S2**. ATAC-seq performed in intestinal epithelial cells from E16.5 *Hnf4αγ^DKO^* versus *Hnf4γ*^+/−^ controls. Genome coordinates for 5,391 accessible chromatin regions become inaccessible upon HNF4 loss (cluster 2 from Fig.6A). Additionally, the full list of HOMER *de novo* motif-calling analysis on these HNF4 chromatin-dependent regions are reported. Finally, the results of GO term enrichment using DAVID analysis for genes with their transcriptional start sites within 20 kb of these HNF4 chromatin-dependent regions are reported. These data correspond to findings in Fig.6.

**Fig. S1.**
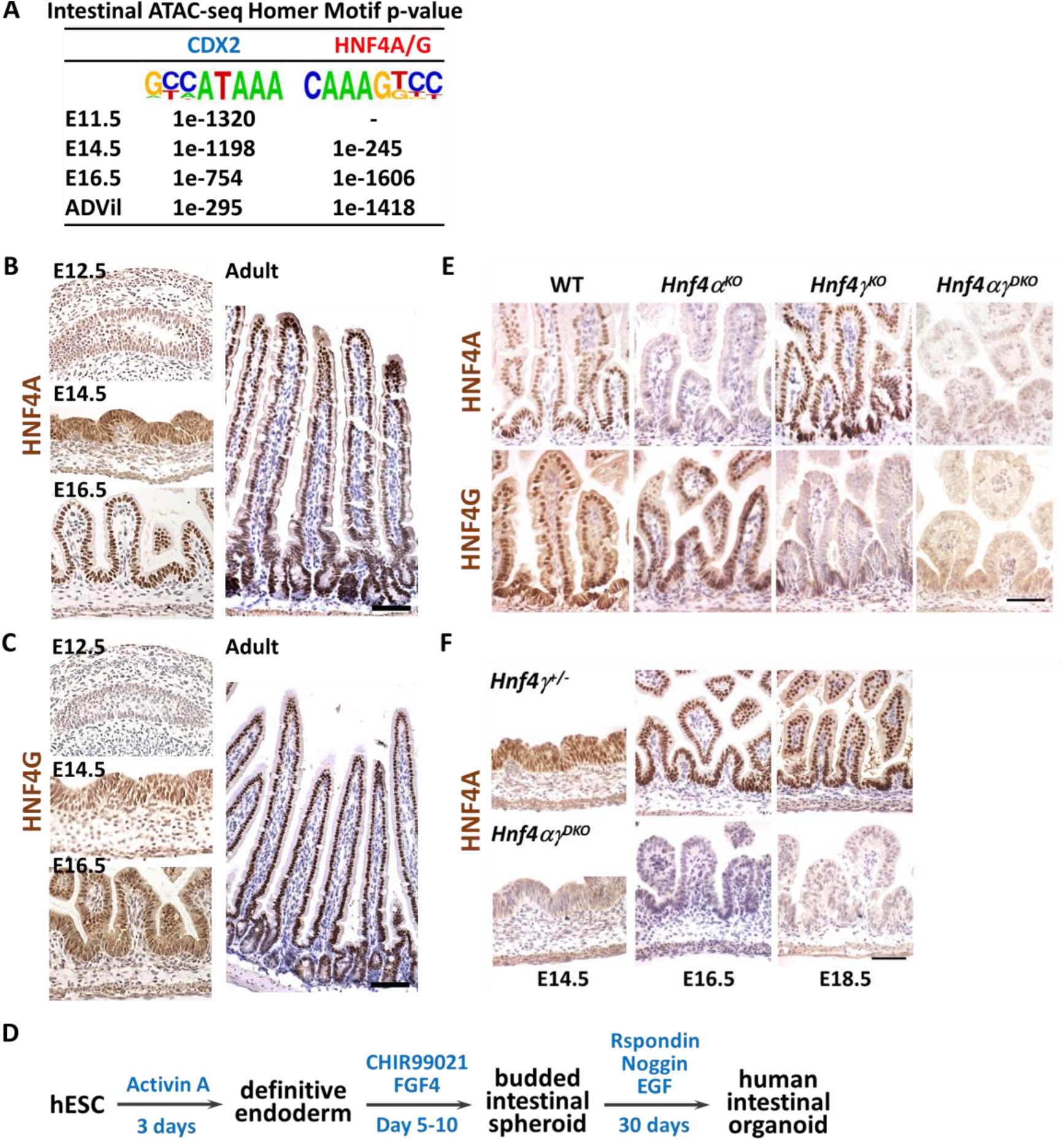
HNF4 expression in the intestinal epithelium across developmental time. (A) HOMER *de novo* analysis of ATAC-seq (GSE115541, n = 2 biological replicates per timepoint, MACS *P* value ≤ 10^−5^) of mouse intestinal epithelial cells at each developmental time shows that CDX2 binding motifs are present as early as E11.5, whereas HNF4 binding motifs are not present until E14.5. HNF4 motifs are increasingly abundant at accessible regions as the intestine matures. Immunostaining of (B) HNF4A and (C) HNF4G shows that relative protein levels of these factors increase with developmental time in mouse (representative of 4 biological replicates, duodenum). (D) Schematic of human intestinal organoids differentiated from hESCs (Tsai et al., 2017). (E) HNF4A and HNF4G immunostaining shows loss of HNF4 in the E18.5 intestinal epithelium of both single and double mutants (representative of 4 biological replicates, E18.5 duodenum). (F) Immunostaining of HNF4A shows loss of HNF4A in the intestinal epithelium of E14.5 *Shh-Cre*+ embryos (representative of 4 biological replicates, duodenum). Scale bars, 50 μm.

**Fig. S2.**
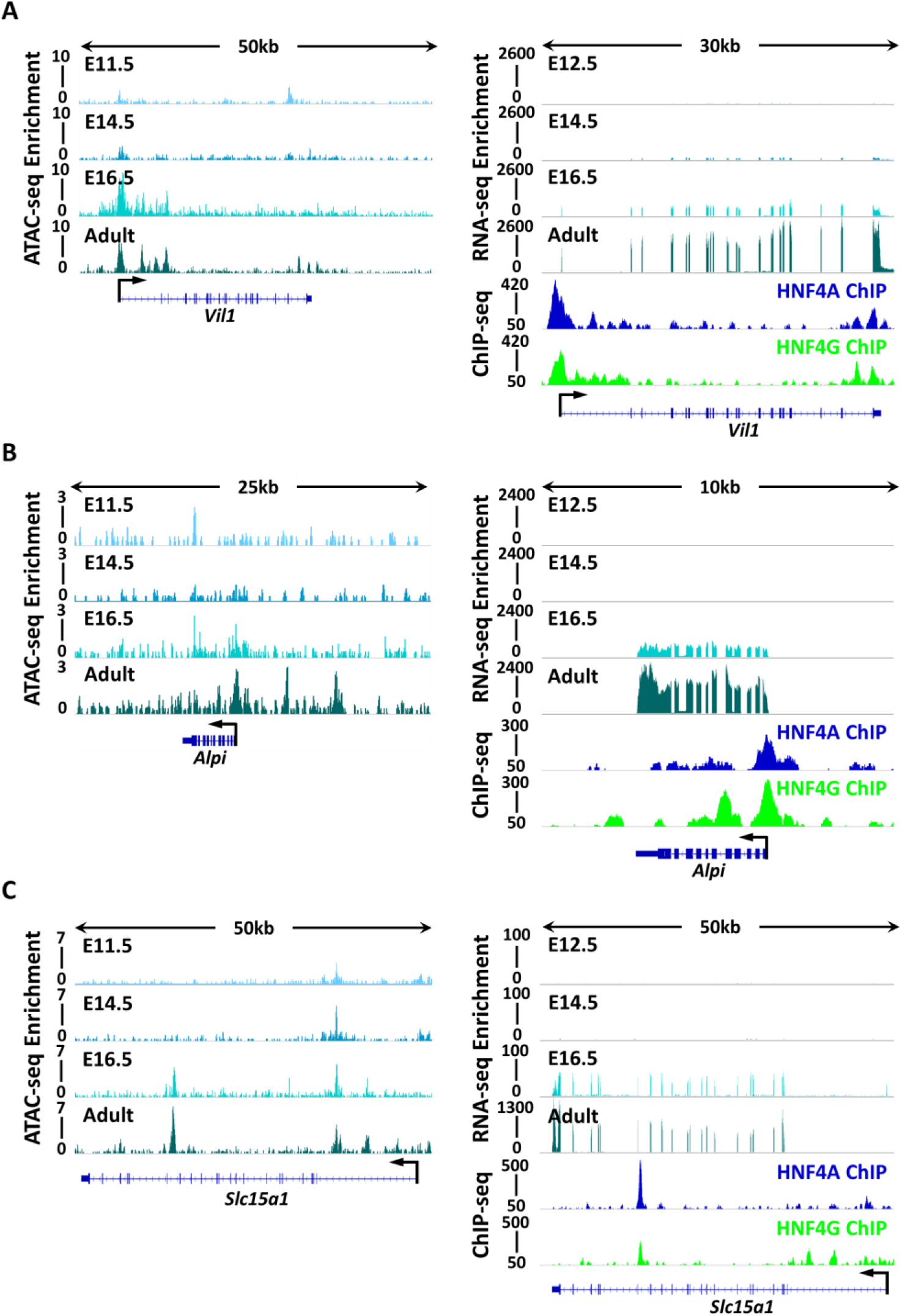
HNF4 paralogs bind to and activate maturation-specific genes. Maturation-specific genes, such as (A) *Vil1*, (B) *Alpi* and (C) *Slc5a1*, show more accessible chromatin (ATAC-seq, left panels) and increased transcript levels (RNA-seq, right top panels) across developmental time, and HNF4 factors bind to the maturation-specific genes in the mature tissue of the adult (ChIP-seq, right bottom panels). n = 2 biological replicates per developmental timepoint for ATAC-seq (GSE115541), RNA-seq (GSE115541) and ChIP-seq (GSE112946).

**Fig. S3.**
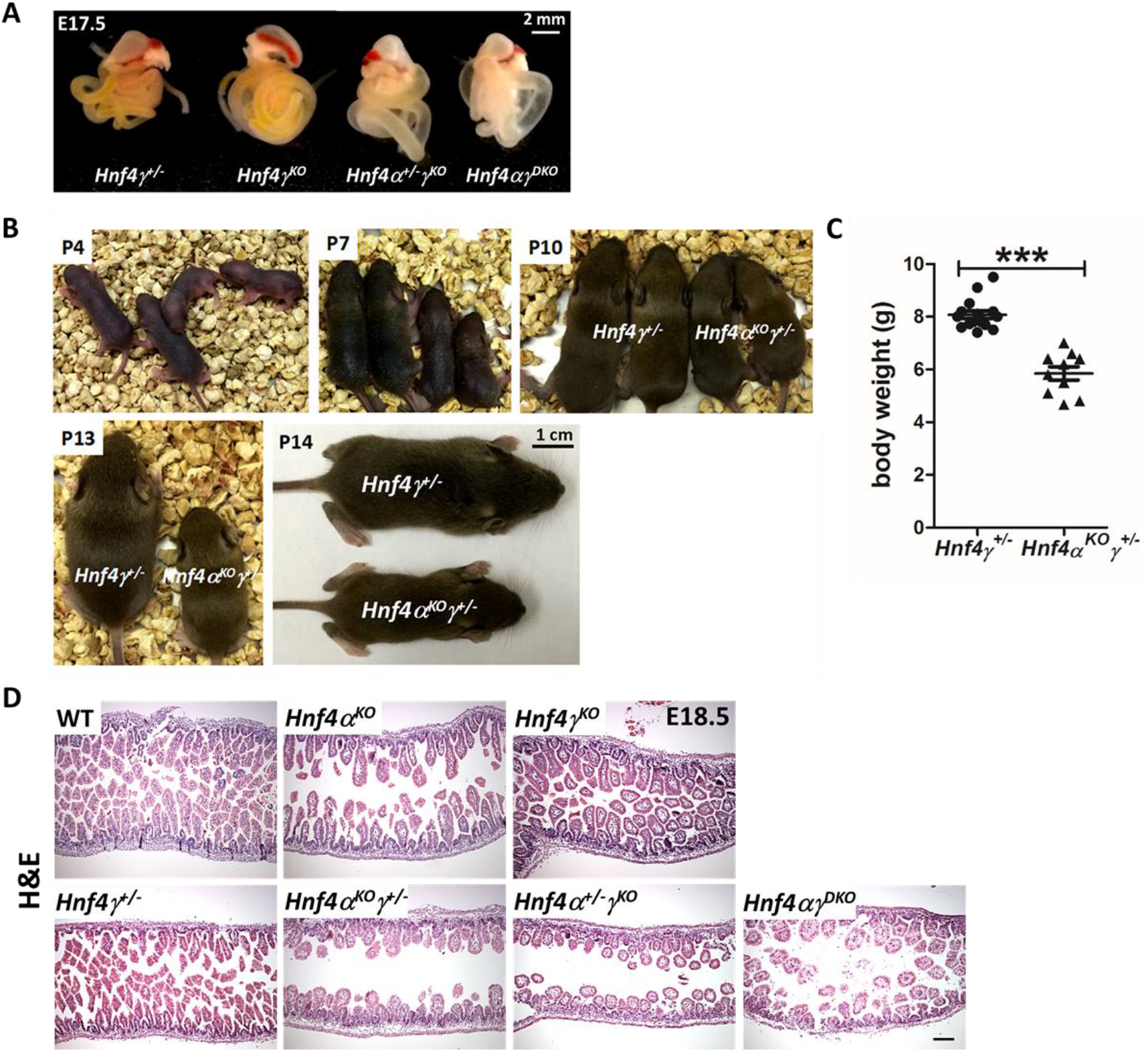
Loss of 3 *Hnf4* alleles in the developing gut leads to growth retardation after birth. (A) Loss of 3 or 4 Hnf4 alleles in developing embryos leads to an underdeveloped intestine with distended and translucent lumen (representative of 4 biological replicates). (B) *Hnf4α^KO^γ*^+/−^ pups can survive after birth but show slower growth when compared to littermate controls. (C) Body weight of P14 pups. Data are presented as mean ± SEM (*Hnf4γ*^+/−^ controls: n = 16; *Hnf4α^KO^γ*^+/−^ mutants: n = 10; Student’s t-test, two-sided at P < 0.001***). *(D) Hnf4* single mutants have a similar morphology to the controls, but loss of 3 or 4 *Hnf4* alleles in developing embryos leads to strikingly stunted villi, as evidenced by H&E staining (representative of 4 biological replicates, E18.5 duodenum; expanded panel from Fig.6H). Scale bars, 50 μm.

**Fig. S4.**
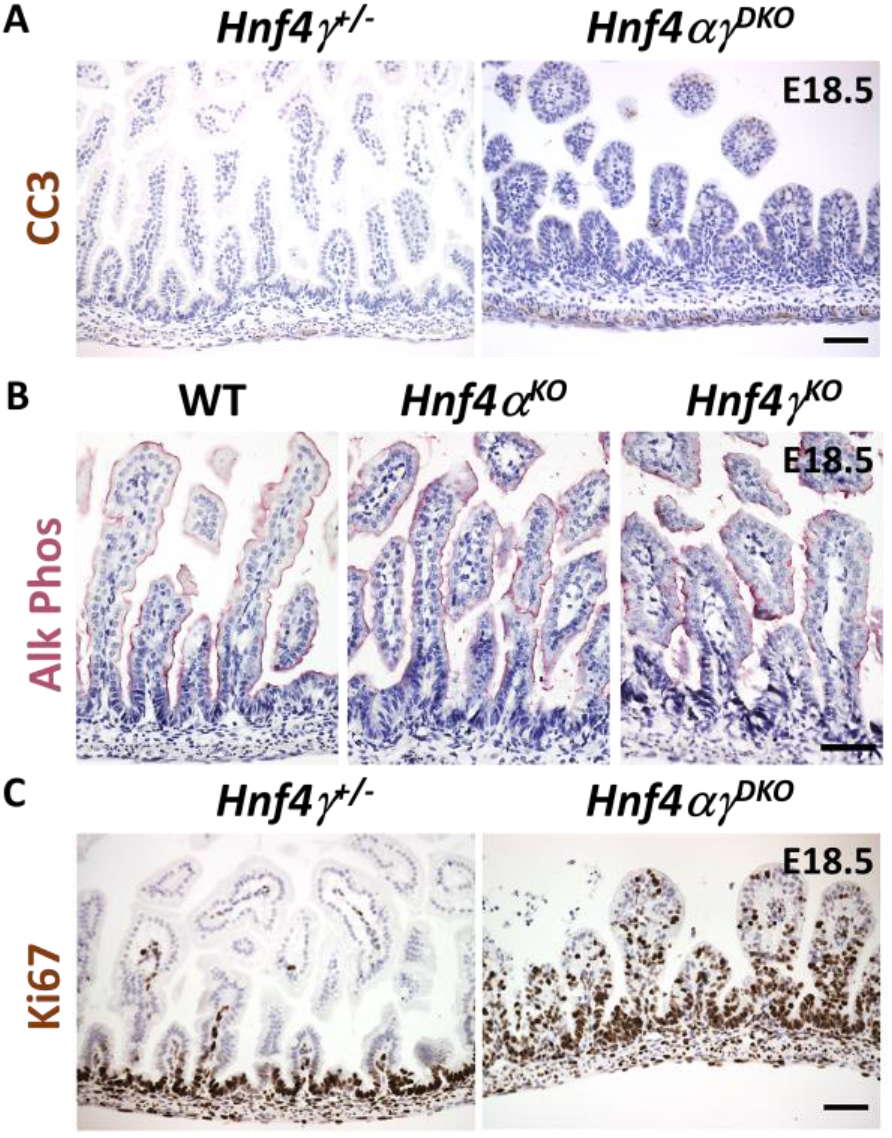
Additional histological and immunochemical features of HNF4 mutant intestines. (A) *HNF4* mutants do not have significant cell death as shown in immunostaining of cleaved caspase 3 (representative of 4 biological replicates, E18.5 duodenum). (B) *Hnf4α^KO^* and *Hnf4γ^KO^* embryos do not have compromised intestinal maturation, as evidenced by alkaline phosphatase staining, indicating redundant roles for HNF4 factors in intestinal development (representative of 4 biological replicates, E18.5 duodenum). (C) Proliferative cells (Ki67+) are observed in the villi of *Hnf4αγ^DKO^*, which may be due to compromised tissue maturation (representative of 4 biological replicates, E18.5 duodenum). Scale bars, 50 μm.

